# Mosaic evolution of a learning and memory circuit in Heliconiini butterflies

**DOI:** 10.1101/2024.04.21.590441

**Authors:** Max S. Farnworth, Theodora Loupasaki, Antoine Couto, Stephen H. Montgomery

## Abstract

A critical function of central neural circuits is to integrate sensory and internal information to cause a behavioural output. Evolution modifies such circuits to generate adaptive change in sensory detection and behaviour, but it remains unclear how selection does so in the context of existing functional and developmental constraints. Here, we explore this question by analysing the evolutionary dynamics of insect mushroom body circuits. Mushroom bodies are constructed from a conserved wiring logic, mainly consisting of Kenyon cells, dopaminergic neurons and mushroom body output neurons. Kenyon cells carry sensory identity signals, which are modified in strength by dopaminergic neurons and carried forward into other brain areas by mushroom body output neurons. Despite the conserved makeup of this circuit, there is huge diversity in mushroom body size and shape across insects. However, an empirical framework of how evolution modifies the function and architecture of this circuit is largely lacking. To address this, we leverage the recent radiation of a Neotropical tribe of butterflies, the Heliconiini (Nymphalidae), which show extensive variation in mushroom body size over comparatively short phylogenetic timescales, linked to specific changes in foraging ecology, life history and cognition. To understand the mechanism by which such an extensive increase in size is accommodated through changes in lobe circuit architecture, we first combined immunostainings of structural markers, neurotransmitters and neural injections to generate, to our knowledge, the most detailed description of a Papilionoidea butterfly mushroom body lobe. We then provide a comparative, quantitative dataset which shows that some Kenyon cell populations expanded with a higher rate than others in *Heliconius*, providing an anatomical parallel to specific shifts in behaviour. Finally, we identified an increase in GABA-ergic feedback neurons essential for non-elemental learning and sparse coding, but conservation in dopaminergic neuron number. Taken together, our results demonstrate mosaic evolution of functionally related neural systems and cell types and identify that evolutionary malleability in an architecturally conserved parallel circuit guides adaptation in cognitive ability.

## Introduction

Brains are the interface between perception and an individual’s response to the environment. How evolution modifies these neural systems to generate adaptive change in sensory detection and behaviour is a key question in evolutionary research [1–3]. On a systems level, neural circuits are the product of often largely conserved developmental programs and operate within the constraints of shared functionality and interdependency [2,4]. These biological relationships can shape how circuits evolve, potentially favouring coordinated evolution across functionally related circuits, or between cell types with shared developmental origins [5]. As functional relationships are determined by connections across macroscopic areas of brains, neural circuits offer the closest brain-wide anatomical correlate to function [6–8]. However, our understanding of what makes some certain circuits more conducive to evolutionary change than others, and which mechanisms are used to enact that change, is still developing [9–11].

By coupling generally conserved cellular components but highly divergent morphologies, insect mushroom bodies, which facilitate learning and memory [12], offer an informative model system to examine the evolutionary dynamics of circuit change. Mushroom bodies are dominated by their main intrinsic cell type, Kenyon cells (KCs) [12–15]. KCs form dense postsynaptic dendritic arborisations on the posterior side of the brain, the calyx, where they receive multisensory input from projection neurons carrying information from primary sensory neuropils. The sparse activation of KCs in response to input from projection neurons represents a widely conserved characteristic of memory circuits, enabling a wide range of information to be encoded in the mushroom body [16,17]. Parallel fibres from the KCs then project anteriorly forming the peduncle, then split into differently distinct sub areas that are defined by their supplying KC types. These are broadly distinguished as α, β, α’, β’ and γ lobes where in most species α and α’ constitute a ‘vertical’ lobe while β, β’ and γ comprise a ‘medial’ lobe [14,18–20]. Each group of neurons connects pre-synaptically to different populations of dopaminergic neurons (DANs) and mushroom body output neurons (MBONs). The resulting circuitry forms the mushroom body lobe mass and is the site of cognitive processes such as learning and memory. Specifically, DANs convey whether an event has positive or negative valence and modify spiking intensity of KCs connecting to MBONs. KCs therefore carry ‘sensory identity’, DANs carry a ‘teaching signal’ of past experience, and MBONs communicate this ‘learned output’ to other circuits in the brain [14]. This simplified wiring logic is, as far as currently known, extremely conserved across insects, and similar architectures have evolved convergently several times across animals [14,21].

However, despite this conserved organisation of cell types, mushroom body lobe anatomy can be vastly different across the insect clade [13,22], particularly in terms of separations between the vertical and medial lobe (S1 Fig). While key insights into lobe morphology of representatives of some groups have been made in recent years [13,14,19,23], a clear evolutionary framework is largely missing, as most comparisons to date have been made between a few species with very deep phylogenetic divergence. This is despite the important role this brain area plays for cognitive processes, and despite a need to contextualise the findings of select model organisms within a wider phylogenetic context.

Here, we aim to understand the evolutionary dynamics of mushroom body circuits, and their capacity to accommodate adaptive change, by leveraging three key characteristics of a Neotropical tribe of butterflies, the Heliconiini. First, Heliconiini comprise a large radiation of closely related species with broadly conserved ecologies [24]. Second, as an established system for eco-evolutionary research, their behaviour and ecology is broadly understood, providing a basis to interpret neuroanatomical data [25]. Third, within this tribe, almost all species of the genus *Heliconius* have a derived suite of traits linked to an ability to exploit a novel amino acid rich food source, pollen, as an adult [26]. Efficient pollen foraging necessitates memorizing sparsely distributed resources across an individually consistent home range to which they show strong site-fidelity [26,27]. *Heliconius* show specific enhancements in certain learning and memory tasks, including increased stability of visual long-term memories and enhanced non-elemental learning [28,29], while performance in most other tasks is unaffected [30]. These specific cognitive demands have driven a 4-fold expansion of the mushroom body volume, relative to the rest of the brain, caused by an even greater increase in KC number and increased visual specialisation to the calyx [28].

The apparent specificity of improvements in a narrow range of behavioural and sensory contexts predicts parallel patterns of mosaic change in the underlying mushroom body circuitry. However, our ability to test this hypothesis has been lacking due to a lack of clarity over how an increase in KC number impacts the broader circuit formed by the mushroom body and its constitutive cell types. We tackle this question by first assessing the structure and interspecific variability in the Heliconiini lobe mass through careful anatomical analysis. This is particularly challenging compared to other species previously studied, as Nymphalid butterflies have spheroid lobes, with no clear split between a vertical and medial portion (S1 Fig). Accompanied by the volumetric expansion in *Heliconius*, the anatomical boundaries of the lobes have previously been obscure [31]. We then provide a comparative, quantitative dataset to analyse changes in expansion rates of all lobe divisions which reflect different KC populations. Finally, we integrate other key cell types, DANs and feedback neurons, by providing qualitative and quantitative analyses of their abundance. Our findings open the door to examine mushroom body circuitry in the light of vast volumetric changes through adaptive evolution to accommodate cognitively relevant behaviour.

## Results and Discussion

### Conserved wiring logic reveals a complex picture of lobe divisions in Heliconiini

To identify homologous lobe divisions reflecting Kenyon cell populations, we used a combination of acetylated tubulin and Horseradish peroxidase (HRP) staining on representatives of Heliconiini with expanded (*Heliconius sp*., approx. 80,000 KCs) and more modest mushroom bodies (in particular *Dryas iulia*, approx. 10,000 KCs). Tubulin stains large parts of the cell, including the axon, thus labelling the axonal tract system that constructs the lobes [32]. HRP is concentrated at the end points of neuronal cells, particularly in synapse-rich areas [33]. Co-labelling against HRP and tubulin therefore offers a comprehensive view of the tract systems and functional domains in the lobes (Fig 1). We further confirmed identities and divisions through Dextran-conjugated dye injections into the mushroom body calyx and Fasciclin-II (FasII) immunostainings (S5 Fig A/B)[34]. Using this approach, we identified three conserved anatomical features used to determine homology [13,14,19,20,23,35]:

**Fig 1:**
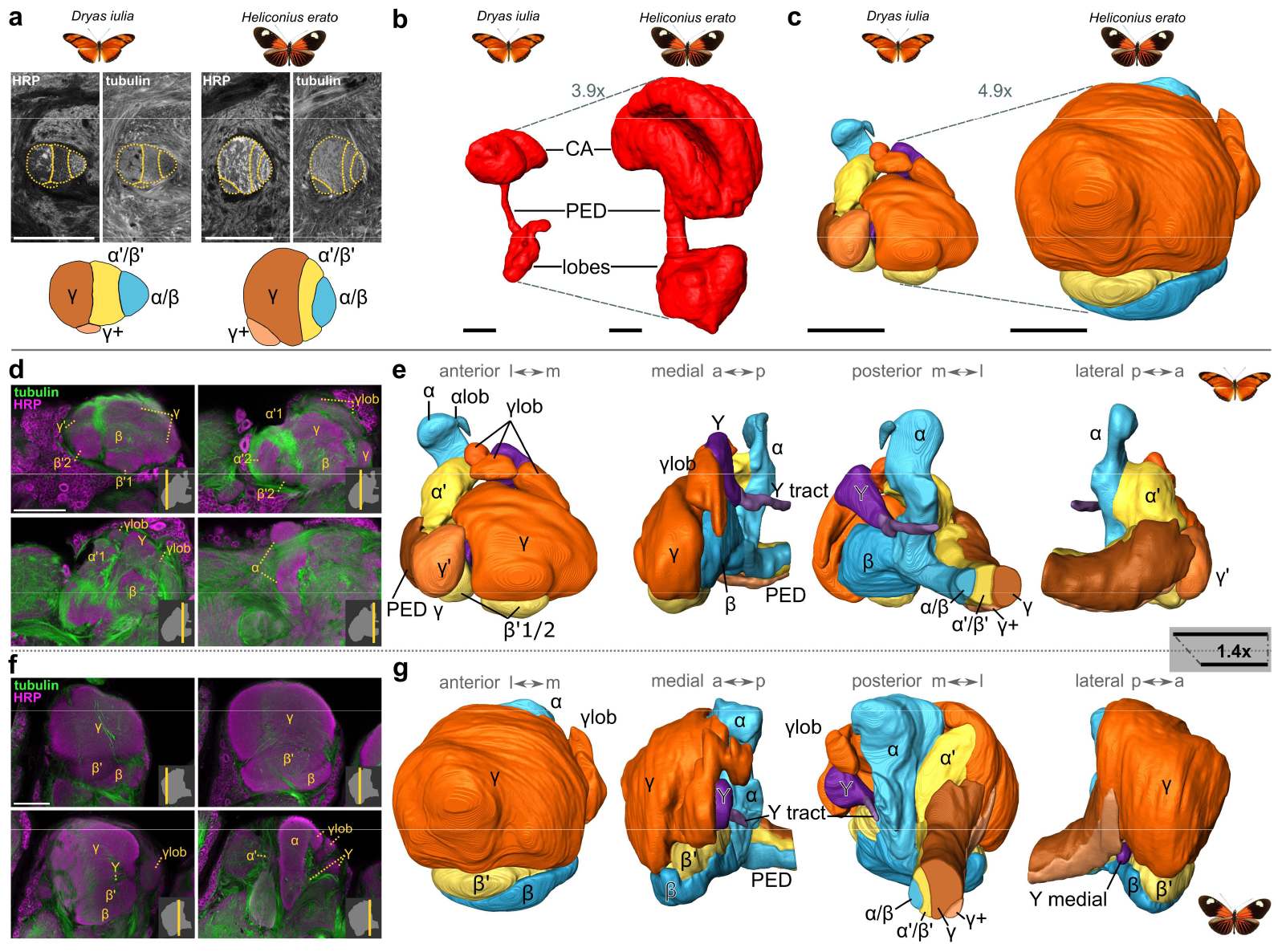
Mushroom body lobe anatomy in Heliconiini and divergence in composition between a non-pollen feeder *Dryas iulia* and a pollen feeder *Heliconius erato*. **a**. Homologous lobe divisions were determined based on the different portions of Kenyon cells running through the peduncle. Shown are single plane sections, with annotations of the different divisions visible, and a coloured reconstitution for both species. **b/c**. The mushroom bodies as a whole are massively expanded in *Heliconius erato* in comparison to *Dryas iulia*, as are the lobes specifically. The fold-change is the absolute volumetric change. **d/f**. Singe plane sections of stainings in *Dryas iulia* (**d**) and *Heliconius erato* (**f**) that illustrate different subdivisions, including denotations of structures and position alongside the anterior-posterior axis in the lower-right corner of each panel; most anterior is top-left, most posterior is lower-right. **e/g**. 3D segmentations in four different orientations with annotations of the structures determined. Scale bars illustrate that e and g are not shown to-scale. See S1/S2 Video for an animated 3D segmentation. All scale bars are 100 μm.

1. The neurites of major KC types are distinguishable inside the peduncle and can be tracked projecting into the lobe mass through differing fibre and synaptic densities (Fig 1A). This means that the subdivisions supplied by KCs are an appropriate proxy for the relative size of the population of KCs.
2. Neurites of these KC populations split at different but stereotypical and conserved locations into their respective subdomains, following the order of γ – α’/β’ – α/β in the peduncle from lateral to medial, and in the lobes from anterior to posterior (Fig 1D-G). The γ lobe, for example, is the most anterior section and supplied by the most lateral peduncle division.
3. Neurites also split into a medial and vertical portion at the end of the peduncle, revealing previously hidden divisions comprising vertical and medial lobes (Fig 1D-G). These lobes are prominently divided by the Y lobe in Heliconiini.

The presence of these three features implies that the wiring logic is conserved between Heliconiini and other insects [13]. These criteria, corroborated by multiple labelling methods (Fig 1, S5 Fig), provide a framework to identify homologous lobe structures. The relative size of each division as well as their shape and relative position, however, differed dramatically in the species examined here (Fig 1). In *Dryas iulia*, which has smaller mushroom bodies, we generally noticed more fine-grained structuring, whereas in *Heliconius* a more simple structural atlas with modified shapes could be identified (Fig 1 D-G, S1/S2 Video, S1 Table), likely due to the massively expanded lobes in this genus compressing anatomical boundaries (see a more detailed description in S1 Text). Throughout Heliconiini we see a large spheroid γ lobe, which in the case of *Heliconius sp*. hides the vertical α lobe from an anterior view. The larger volumes of lobe divisions, particularly the γ lobe, in *Heliconius* seem to have led to a flattening of β and β’, located ventrally to the dominant γ lobe, while they are relatively spheroid in *Dryas iulia* and located posteriorly to the γ lobe. In contrast, the Y lobe and tract are largely conserved across Heliconiini. We also noticed previously unmentioned aspects common across Heliconiini, such as an additional KC portion we describe as ‘γ+’ innervating the medially positioned γ lobelets (S1 Text, Fig 1E/G, S5 Fig A/B), as well as additional divisions, directly visible only in *Dryas iulia*, which likely represent specific subtypes of KCs, also characterised by different portions of DANs and MBONs [14,36,37].

To our knowledge, these analyses provide the most detailed identification of lobe divisions and KC populations in Papilionoidea (butterflies), and reveal high degrees of complexity but a conserved wiring logic behind the spheroid lobes of Heliconiini and Nymphalids generally [38]. Such shape differences are not surprising considering the vast differences in KC number across insects: approximately 2,000 KCs per hemisphere in the moth *Spodoptera* and the fly *Drosophila* versus 10,000 and 80,000 per hemisphere in non-pollen feeding and pollen-feeding Heliconiini, respectively [14,19,28]. Additional comparisons of the larval mushroom body revealed a structure which more closely resembles those of *Drosophila* larval mushroom bodies [39,40] (S2 Fig), with no obvious differences between Helconiini genera (S2 Fig), which implies that all differences present in the adult must have arisen during late larval and pupal development (see also S1 Text).

### Mosaic expansion of lobe divisions indicates malleability in lobe sub-circuits

We next employed several statistical methods to provide a comparative framework to test the hypotheses that a) there are species and clade-specific effects of expansion in the lobes and b) that these occurred largely across all lobe divisions, illustrating functional and developmental dependency between them (all statistical results can be found in S2 Table). For this, we generated a dataset of FasII and HRP stained brains, with which we measured the size of the identifiable lobe divisions across both sexes of four species (*Dione juno, Dryas iulia, Heliconius erato* and *Heliconius melpomene;* Fig 2A), in both young and aged butterflies (S3 Fig). We found minimal effects of sex and age (S3 Fig, S1 Text for more information) and therefore focus our discussion on interspecific variation in young animals.

**Fig 2:**
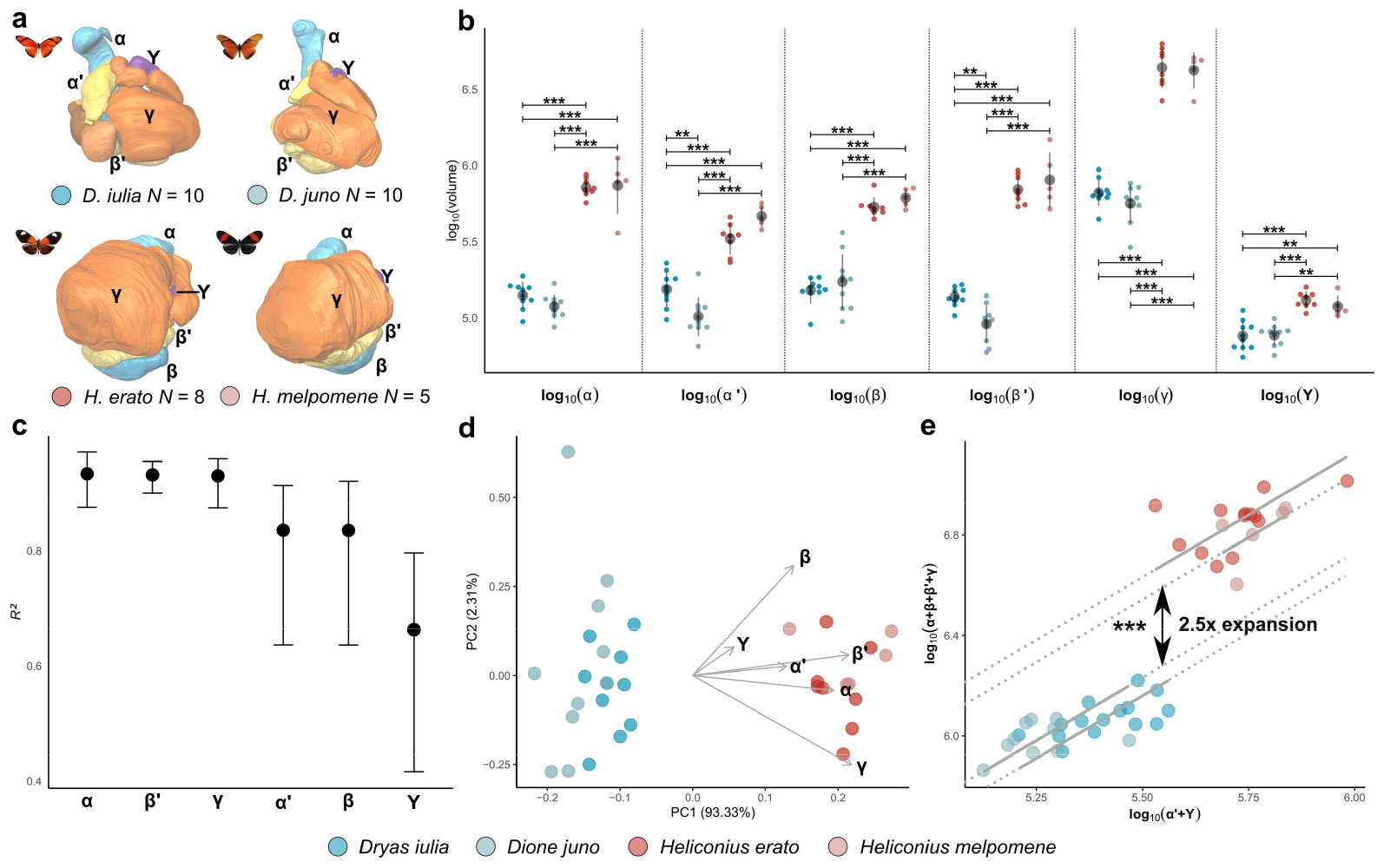
Statistical analysis of mushroom body lobes across four Heliconiini species reveals diverging patterns of expansion. **a**. Data used, showing representative 3D segmentations for each species and number of individuals included. **b**. Volume of each lobe division across species, with indications of significance based on post-hoc pair-wise tests. **c**. *R*^*2*^ across lobe divisions to illustrate extent of expansion between outgroup and *Heliconius* species. **d**. Principal Component (PC) analysis of the volume of all structures to examine overall variation and determine global patterns of difference per structure. Indicated are loading directions per division. **e**. Summary of *smatr* analyses (see also S4 Fig) of scaling relationships between lobe divisions. Structures that are relatively more expanded than others were grouped and contrasted to those that are relatively less expanded. Indicated are significant elevation differences between *Heliconius* and both outgroup species (species-specific differences are reported in the data file). This confirmatory analysis illustrates higher rates of expansion in specific lobe divisions and Kenyon cell populations.

Our statistical analyses revealed species as a better predictor for variability in lobe subdivision size than clade (*Heliconius* vs. not-*Heliconius*) (*X*^*2*^ = 43.142, *P* < 0.001) suggesting a degree of variability not explained by mushroom body expansion alone. Although lobe structure volumes vary significantly across species (in comparison to the null model; *X*^*2*^ = 189.602, *P* < 0.001), importantly, the largest effects were seen between *Heliconius* and non-pollen feeding outgroups. This was corroborated by *post hoc* pair-wise tests examining species effects in each structure separately (Fig 2B). In all cases, both *Heliconius* species had significantly larger lobe subdivisions than both the outgroups, but with additional variation between the α’ and β’ lobes being smaller in *Dione juno* than in *Dryas iulia* (S2 Table). This was expected given the absolute size differences in lobes and mushroom bodies overall [28].

However, an examination of effect sizes (Fig 2B/C) reveals variation in the extent of expansion across lobe structures. We found that while species identity across all structures explained considerable amounts of variation (66.3-93.4%), over 90% of variation differences in α, β’ and γ lobes was assigned to species, but species had less explanatory power for the α’, β and Y lobes. Moreover, the variation of *R*^*2*^ itself was much lower in α, β’ and γ than in the other three structures. A multivariate analysis (PCA) also confirmed clear separation between clades (Fig 2D) but the values and orientation of PC loadings indicated one grouping of α, β’ and γ lobes, and a second grouping of α’, Y and β with a different orientation, mirroring patterns in effect sizes.

To corroborate this evidence of inter-lobe variability, we performed pair-wise allometric scaling analysis between all pairwise combinations of lobes [41](Fig 2E, S4 Fig). These tests identify conserved or inconsistent (non-allometric) scaling, revealing any discrepancy in size variation. These analyses revealed that the Y lobe and α’ lobes are consistently less expanded in *Heliconius* species in comparison to all other divisions (insets and significant elevation differences – i.e. clade shifts –indicated in S4 Fig). In contrast, the α, β, β’ and γ lobes show greater expansion in *Heliconius* species, relative to the Y and α’ lobes, but are more consistent when compared to each other. Grouping these two categories of lobe divisions together (i.e. [α, β, β’, γ] and [Y, α’], confirms non-allometric scaling between them, with a significant shift in the y-axis intercept (a grade-shift) (Fig 2E, *W* = 100.526, *P* < 0.001) indicating a 2.5 fold expansion of α, β, β’ and γ over α’ and Y populations in *Heliconius*.

In summary, we find that while all lobes are expanded in *Heliconius* compared to outgroup genera, groups of lobe subdivisions show variability in the degree of expansion, with categories of lobes co-expanding differentially. This differential expansion of lobe structures and KC populations illustrates a degree of evolutionary ‘malleability’ in the internal mushroom body circuit. KCs are typically generated from cell lineages derived from two or four neuroblasts [39,42], with KC subtypes produced sequentially through changing patterns of transcription factors expression, producing a temporal cascade that generates neuron diversity [39,43]. Hence, during mushroom body expansion in *Heliconius*, neurogenesis has been altered not only to produce more KCs, but to alter the relative timings of KC subtypes. Changes in the relative amounts of certain KC types have the potential to shift the representation of sensory information these subgroups receive [28]. Indeed, *Heliconius* calyces show a pronounced shift in the volume receiving visual input, and our data may suggest such a large shift in visual projection may be represented by shifts in specific KC types. In addition, lobe divisions can be traced to calyx substructures because concentric enwrapping of KC populations in the calyx is modified into a layered organisation in the peduncle which then extends into the lobes [44]. We were able to observe a similar pattern and rotation in our data (S5 Fig A’’’). Hence, this morphological correspondence allows us to hypothesize a correspondence in function between visual calyx and lobe structures, with the most outer calycal layer, in *Heliconius* the region receiving visual input, corresponding to the most lateral portion of the peduncle and most anterior portion in the lobes, incidentally the portion that constructs the massively expanded γ lobes.

Importantly, different populations of KCs concentrated in specialised lobe regions have been assigned to specific cognitive functions. The α/β lobes have consistently been associated with long-term memory formation [45–49], while the γ lobes have been connected to more short-lived memory traces [50,51]. Currently, one model of memory formation involves a memory trace being first formed in γ neurons, then corroborated in α/β neurons over longer periods of time [52–54]. In *Drosophila*, the γ lobe has also been strongly associated with visual memory formation [55,56], despite visual input accounting for only 8% of its innervations [14], while in ants the vertical lobe (α, α’ and potentially a γ’) is strongly implicated in visual memory and visually guided navigation [57]. The particular expansion of key components of the long-term memory trace system – α, β, γ lobes – together with a volumetrically dominant γ lobe, may align closely with evidence of specific improvements in long-term visual memory in *Heliconius* [28,29].

### A conserved set of cell groups innervates the lobes with expanded numbers of GABA-ergic feedback neurons in *Heliconius*, but conservation in numbers of DANs

Ultimately, any expanded populations of KCs need to connect to corresponding cell types that regulate and carry forward any valence signals. To understand components of the circuitry innervating the mushroom body lobes besides the KCs, we performed a series of neurotransmitter stainings using conserved antigens to examine GABA (γ-Aminobutyric acid)-ergic neurons (Fig 3B), dopaminergic neurons (Fig 3D) and serotonergic neurons (S5 Fig C). We corroborated results using Dextran injections into the mushroom body calyx as well as FasII antibody stainings (Fig S5 A and B). Available antibodies for other labels (particularly Glutamate and Acetylcholine) were not immunoreactive in our species (S1 Table).

**Fig 3:**
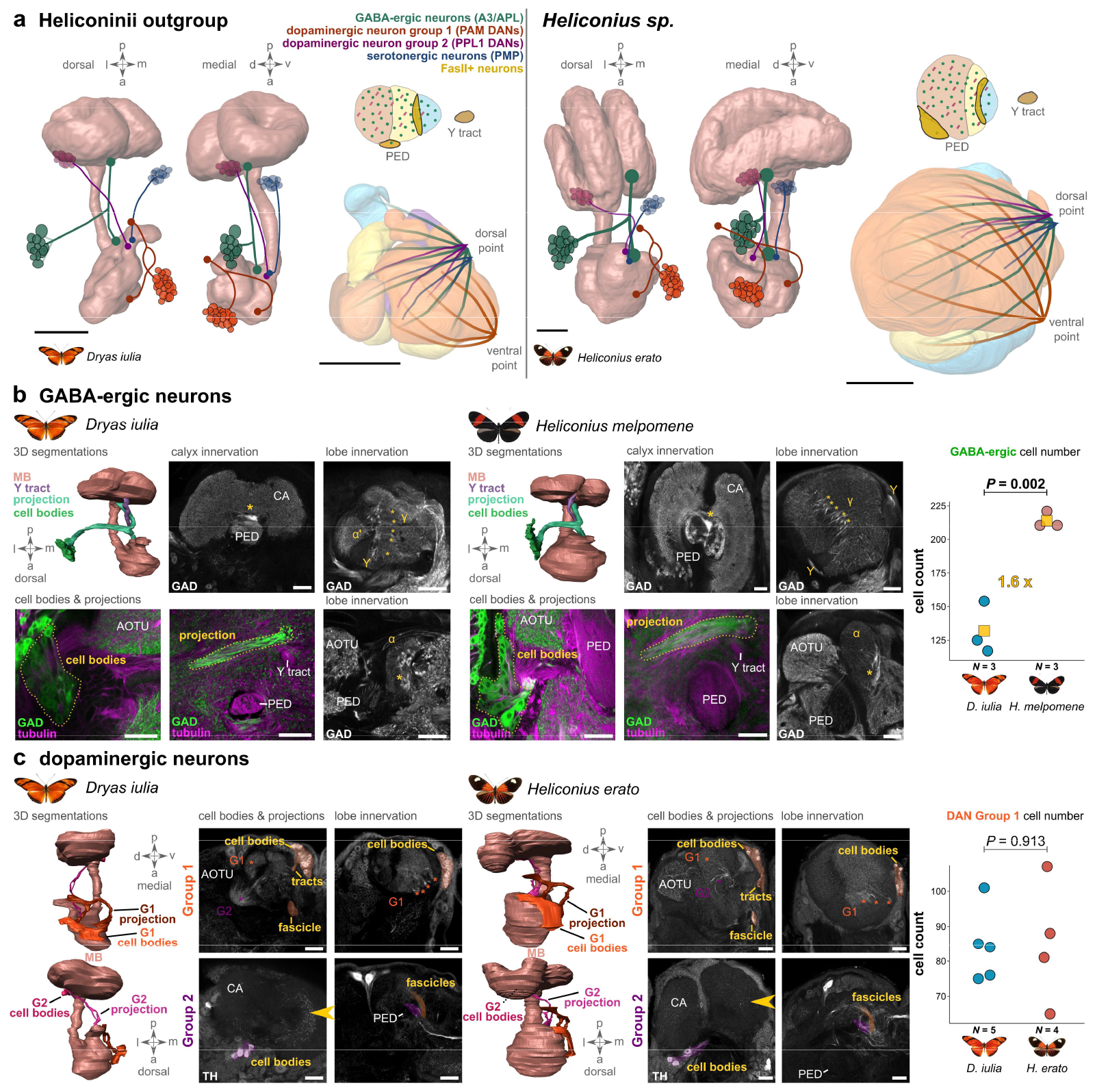
Analysis of non-Kenyon cell groups supplying projections into the mushroom body lobes across outgroups and *Heliconius* species. **a**. A summary of the results in b, c and S5 Fig. First, the approximate position of analysed cell groups are shown in two orientations across the whole mushroom body. Second, the innervation patterns inside the peduncle, Y tract and innervation pattern inside the lobes are shown. Note that most broadly, there were two fascicle entry points into the lobes, a dorsal and a ventral point. Cell group location and projection pattern were generally conserved, but cell number diverges for some cell groups. 3D segmentations are from identical sources than in Fig 1. **b**. Analysis of GABA-ergic neurons using the antibody targeting glutamic acid decarboxylase (GAD) revealed feedback neurons innervating the calyx and lobes simultaneously. While conservation occurred anatomically, cell numbers were 1.6-fold higher in *Heliconius*. **d**. Analysis of dopaminergic neurons (DANs) using an antibody targeting Tyrosine Hydroxylase (TH) revealed two prominent cell groups innervating the lobes. Group 1 is a large cell group with conserved neuron numbers across species, innervating into the ventral point of the lobes through a tract that bends across the ellipsoid body. Group 2 was a smaller group of cells innervating through the dorsal point. All scale bars in a are 100 μm and in b/c 50 μm.

We detected conserved patterns of cell group locations, projections and innervations (collated in Fig 3A). We identified a dorsally located conflation point of several projections (GABA-ergic, serotonergic and one of two dopaminergic neurons) and one prominent ventral point where a large group of dopaminergic neurons passes into the lobes (Fig 3 A/B). Within the peduncle and Y tract we also observed dot-like innervations in GABA-ergic and dopaminergic neurons, while FasII labelling revealed a distinct peduncle layer as well as the γ+ and Y portion. Given the functional importance of GABA as a general neurotransmitter and one of three that are characteristic of MBONs and the importance of dopaminergic neurons in carrying signals of reward or punishment [14], we examined GABA-ergic and dopaminergic neurons more closely.

We examined DANs using a Tyrosine Hydroxylase (TH) antibody (S1 Table). DANs are a major part in the learning circuit as they carry information about reward and punishment, innervating specific regions of KC axons to modify signal strength that is then projected forward by MBONs [12]. Our labelling revealed two cell groups that showed highly conserved cell body locations, projection patterns and innervation. Cell group 1 has cell bodies anterior lateral to the lobes, nestled directly next to the neuropil with a projection joining a major fascicle that projects first ventrally to the ellipsoid body of the central complex and then bends dorso-posterior. The projection splits with one half reaching into the superior medial protocerebrum (potentially pre-synaptically) (lower-right panel in Fig 3C), and the other half reaching the lobes (upper-right panel; potentially post-synaptically). As such, Cell group 1 shows close correspondence to PAM DANs identified in *Drosophila* [58,59], thought to convey reward signals [12,14], while the projection seems very similar to the SEC (supraellipsoid commissure) identified in *Tribolium* [32]. Cell group 2, much lower in number (in both species approximately 10) but with large soma sizes, originates ventro-laterally to the calyx, then projects along the anterior-posterior axis before bending ventrally into the lobes. In contrast to group 1, which is the only cell group examined here that enters the lobes through a ventral conflation point, it projects into the lobes dorsally. This cell group most likely corresponds to the second major class of DANs, the PPL1 neurons [58,59] which convey punishment [12,14]. A similar projection pattern, but from a ventro-medial position to the calyx was detected in serotonergic neurons, with very sparse labelling inside the lobes through the dorsal conflation point (Fig 3A, S5 Fig C), most likely corresponding to PMP neurons identified in *Drosophila* [58,60].

Importantly, PAM DANs (group 1) show no change in cell number across representatives of *Heliconius* and *Dryas* (*t* = -0.113, *P* = 0.913, Fig 3C, manual counting). We also noticed sparse labelling inside the calyx (potentially stemming from the spot-like TH labelling we identified in the peduncle (Fig 3A)), where in both species the inner ring of calyces was innervated, which may overlap with the olfactory region of the calyx in *Dryas* [28]. Intensity was much reduced in *Heliconius*, which in light of the conserved DAN numbers but massive KC increase might stem from a dilution effect. Nevertheless, innervation of calyces brings further indication of unconventional innervation patterns with afferents reaching calyces through other cell types than projection neurons.

We next examined GABA-ergic neurons through anti-GAD (glutamic acid decarboxylase) immunolabelling (Fig 3B) [14] and detected a prominent cell group anterior-lateral to the lobes, in the crevice between optic lobe and central brain that projects posterior-medially at first, in a thick fibre bundle perpendicular to the peduncle, then splits medially into a tract that enters the calyx just dorsally to the peduncle and an anterior part that innervates the lobes globally through the dorsal conflation point (this pattern was corroborated through injections that revealed this cell group again, S5 Fig A’/A’’’’’). While the overall pattern of projections was highly conserved, cell numbers (determined through manual counting) were significantly higher in *Heliconius melpomene* than in *Dryas iulia* (*t* = 6.947, *P* = 0.002, Fig 3B), a pattern that was visibly consistent across other species of Heliconiini.

The projection pattern of this GABA-ergic cell group is consistent with GABA-ergic inhibitory feedback neurons [61,62], in *Drosophila* termed anterior paired lateral (APL) neurons [14,16,37,63] and in bees called A3 feedback neurons [64,65] (note that our labelling most likely includes A3v as well as A3d neurons). They are often characterised as MB extrinsic neurons, but definitions of MBONs would allow them to be characterized as MBONs as well. Nevertheless, projection patterns of these feedback neurons are conserved with our data on lepidopteran species. Interestingly, these inhibitory feedback neurons have been found to be responsible for sparse coding of KC signals, which is essential to generate precise combinatorial signalling of stimulus identity inside the mushroom bodies [16,63,66]. The increase of such feedback neurons in species with more KCs may reflect the need to adjust mechanisms to generate sparse coding [64]. Moreover, GABA-ergic feedback neurons have been identified as necessary for non-elemental and reversal learning in bees [67,68]. Particularly with regards to non-elemental learning, increases in cell number in these GABA-ergic feedback neurons neatly parallels previous cognitive data across Heliconiini, where non-elemental learning in *Heliconius* is enhanced relative to outgroup Heliconiini [30]. Our results offer a quantified insight into the evolution of this cell group in this cognitive context. While Devaud et al [67] highlight important differences between the cell groups in bees and *Drosophila*, to our knowledge, these feedback neurons have not been broadly addressed in terms of their evolution as well as in a visually guided context until now.

In summary, mirroring a mosaic change in KC populations, we see a mosaic change of major cell types closely integrated into the same circuit, namely a conservation in DAN numbers, but an increase in GABA-ergic feedback neurons. Conserved DAN numbers may suggest that regulation by DANs of sparse signalling being projected by the many more KCs might not need to occur in *Heliconius*, but for adequate sparse signalling to occur in the first instance a greater number of feedback neurons is required.

## Conclusion

In this work, we have leveraged a phylogenetically recent, but extensive shift in brain composition which is associated with derived patterns of learning and memory, to understand how substantial volumetric differences in a key brain area are underpinned by changes in cell populations. In doing so, we provide an anatomically rich, quantitative analysis of mushroom body lobe anatomy, and discuss the fascinating circuitry of the mushroom body lobes in a clear evolutionary framework. This revealed a complex anatomical picture with substantial differences to other insects – including potential novelties, but one that is still based on a conserved wiring logic in the mushroom bodies. We have demonstrated that KC sub-populations were expanded to differing extents during the evolution of pollen feeding in *Heliconius*, illustrating a mosaic pattern of neural evolution predicted by the specificity of behavioural differences observed between *Heliconius* and other Heliconiini. This is indicative of an evolutionary malleability of neural circuitry and particularly illustrates that volumetric change in brain area sizes, even when highly localised, is unlikely to occur without concomitant changes in other parts of their broader circuit.

An important assumption we have taken throughout this work is that lobe divisions and their size are an adequate proxy for the size of the KC populations they are made of, i.e. with an expanded functional domain should also come an expanded population of KCs. While this might also stem from increased numbers in MBONs and DANs (Fig 3), the most pronounced and absolute difference in magnitude are differences in KC number (approx. 10,000 per hemisphere in *Dryas iulia*, an outgroup species, versus 80,000 in *Heliconius melpomene*) [28]. DANs did not show changes in cell number, and while feedback neuron increases could explain some expansion in lobe divisions, these GABA-ergic neurons we identified only showed a 1.6x increase, involving a much smaller number of cells (100-250 per hemisphere). Moreover, the shifts in proportions of KC axon groupings inside the peduncle (Fig 1A) correspond to the expansion patterns we determined statistically inside the mushroom body lobes. Together, these data therefore strongly imply that the main contributing factor to differential lobe expansion are the underlying KC populations.

More broadly our data are consistent with models of behavioural change that are brought about through localised replication of cell types within conserved circuits [69], with differential expansion and contraction of sub-circuits reflecting the relative importance to the behavioural phenotype under divergent selection. Critically, the mosaic evolution of lobes and cell types we observe highlights the evolutionary malleability of insect learning and memory circuits on a relatively shallow phylogenetic scale, while also emphasising the importance of co-evolution among functionally dependent cell classes, key predictions of evolutionary models of adaptive brain structure [2,5].

## Material and Methods

### Animal husbandry

Heliconiini butterflies were ordered as pupae from commercial suppliers (The Entomologist; https://butterflypupae.com/ or a Costa Rica Entomological Supply; www.butterflyfarm.co.cr). Upon arrival, they were attached posteriorly to a microfiber cloth in a pop-up cage. Upon eclosion, they were given individual IDs, marked on their forewings, to later identify their age when sampling. They were housed in 2 x 2 x 2 m mesh cages at 26°C, 80% humidity and a 16 h/8 h light/dark cycle, with *ad libitum* feeding that consisted of flowering *Lantana* plants as pollen and nectar supply and artificial feed consisting of 20% sugar and 5% critical care formula (VETARK, Winchester, UK) in water. To obtain larval brains, we supplied *Heliconius erato* and *Dryas iulia* with *Passiflora biflora* as a host plant. Eggs were removed daily and larvae were reared in plastic pots, supplied with *Passiflora* leaves. We selected larval stage 2 (L2) larvae, 24 h after hatching, with darkened cuticula and larger head capsule, distinguishing them from the short stage 1 (L1) phase.

### Datasets and data availability

We used several datasets with different groups of species and stainings for specific purposes. All described patterns in Fig 1, 3 and S5 Fig were tested for consistency across a dataset of at least three individuals per species. In the case of larval brains (S2 Fig) we used a dataset across several L2 as well as L1 and L3 stages to verify morphologies, and we observed strong conservation of the general pattern we present above. For all statistical analyses, besides testing for age effects, we used a dataset of naïve animals, dissected less than 24 h after eclosion. This provided the clearest view of the mushroom body lobes, which undergo significant post-eclosion growth, further obscuring anatomical boundaries [31]. As post-eclosion growth does not involve adult neurogenesis [70], the morphology of the young adult lobes will reliably reflect variation in KC sub-types. Nevertheless, we tested for age effects (S3 Fig), using *Dryas iulia* and *Heliconius erato* aged to between 9-10 days old, at which point they are sexually mature, to assess for differential effects of age on relative lobe size (see statistical analysis).

All datasets used for images (the processed image file and 3D segmentation) as well as the complete datasets of volumes for the statistical analysis, including a script, are supplied at https://tinyurl.com/5n7csbmy [71]. All datasets for statistical analyses as well as the results are supplied in S2 Table.

### Dissection, fixation and immunostaining

All procedures on adult specimens were highly similar to published protocols [28,72]. Adult butterflies were cold-anesthetized for a few minutes and then decapitated. Antennae, proboscis and palps were removed and the head was pinned with anterior to the top. Dissection and fixation of adult brains in Heliconiini butterflies followed three main steps. 1. Opening of the head to reveal a window to the brain and provide exposure to fixative *in situ*, 2. fixation and 3. removal of the brain out of the head capsule post-fixation. Step 1 was performed in HBS (HEPES-buffered saline; 150 mM NaCl; 5 mM KCl; 5 mM CaCl_2_; 25 mM sucrose; 10 mM HEPES) and by cutting two slits along the edge of the eye and head cuticle. Thick cuticle stalks behind the proboscis were cut, the head cuticle was lifted and tissue cut close to the cuticle, including the antennal nerves. In step 3, after fixation, the eye cuticle and eye tissue were carefully removed. All trachea on the anterior side were removed before the brain was lifted out of the head capsule and the posterior trachea carefully removed.

Larval brains were dissected in HBS by first removing most of the abdomen of the larva, then teasing apart the dorsal cuticle to expose the ventrally situated ventral nerve chord (VNC). With the VNC as guidance, the brain was further exposed. From anteriorly, the mouth parts were torn and teased apart to expose the brain fully. It was then lifted from the remaining body cavity. Where possible, several VNC ganglia were retained to ease handling of the small tissue.

We used different fixatives and durations for different immunostainings and tested various antibodies. All tested conditions, including unsuccessful antibody stainings, are reported in S1 Table. In all staining data reported here we used 1% Zinc-Formaldehyde (ZnFA; 18.4 mM ZnCl_2_, 135 mM NaCl, 35 mM sucrose, 1% formalin) as described previously [72]. We fixed brains for approximately 24 h at room temperature on an orbital shaker. We fixed larval brains in the same solution for 4 h.

After fixation and dissection, all brains were subjected to a two hour incubation in Dent’s solution (80% Methanol, 20% DMSO) followed by a rinse and subsequent -20°C storage in methanol. We then rehydrated them in a descending methanol series (90%, 70%, 50%, 30%) diluted in 0.1 M Tris-HCl (pH = 7.4), each step as a 10 min wash, followed by a wash in 100% 0.1 M Tris-HCl.

Sectioning was performed for all adult brains to allow deeper and thorough antibody penetration, high quality imaging and ease of analysis. For this, we first embedded the rehydrated brains in 5% low-melting point Agarose (#16520-050, ThermoFisher Scientific, MA, USA) and fixed them on a magnetic disk holder using cyanoacrylate glue, maintaining a non-tilted anterior-posterior vertical axis as much as possible and having cut one corner into the gel to be able to later correctly identify orientation. We then used the Leica Vibratome VT1000-S (Leica Biosystems, IL, USA) to section at 80 μm with a speed and frequency of 5.

We subsequently rinsed with PBS-d (0.1M PBS [BR0014G, ThermoFisher Scientific, MA, USA] with 1% DMSO), performed a 30 minute permeabilization wash with permeabilization buffer (PBS-d and 2% Triton-X-100), rinsed once again with PBS-d and then proceeded to blocking using 5% of NGS (normal goat serum, G9023, MERCK, Germany) in PBS-d for 2-4 hours. We then applied the first antibodies with the appropriate dilution (see S1 Table) in blocking buffer and incubated them for three days at 4°C on an orbital shaker. Antibody incubation was followed by one rinse and three 30 minute washes with PBS-d. Secondary antibodies were diluted in blocking buffer (see S1 Table) and incubated for three days at 4°C on an orbital shaker. This was followed by a rinse in PBS-d and a 45 minute incubation of DAPI (1:1000) in water with 0.2% Triton-X-100. This was followed again by a PBS-d rinse and four 10 minute washes. Then we washed brains for an hour in PBS, transferred them to 60% glycerol in PBS overnight for approximately 12-16 hours at 4°C or 2-4 hours at room temperature. Brains were then transferred to 80% glycerol for an hour and mounted on frosted object slides (J1800AMNZ, ThermoScientific, MA, USA) covered with #1.5 size coverslips and sealed with nail polish.

L2 larval brains were treated identically, besides a rinse in PBS-d following rehydration and then proceeding to blocking procedures. Before incubation in 60% glycerol, however, we washed them in 30% glycerol/PBS for 1 h.

### Dextran injections into the mushroom body calyx

To label and identify pathways from Kenyon cells to other brain areas and to potentially identify new cell groups, we injected dextran into the mushroom body calyces. These injections followed procedures described elsewhere [28]. In short, butterflies were cold-anesthetized on ice before being secured in custom-made holders with a plastic collar around the cervix. A waterproof barrier was then created using low melting point wax around the head to prevent leakage of a ringer solution (composition: 150 mM NaCl, 3 mM CaCl_2_, 3 mM KCl, 2 mM MgCl_2_, 10 mM HEPES, 5 mM Glucose, 20 mM Sucrose). Subsequently, a portion of the posterior head cuticle was removed to expose the dorsal region of the brain under the ringer solution. The tracheal covering over the calyces was carefully removed, and under filtered illumination, the tip of a pulled glass capillary (G100-4, Warner Instruments, CT, USA), loaded with crystals of fluoro-ruby (dextran-tetramethylrhodamine: 10,000 MW, D1817, Thermo Fisher Scientific, MS, USA) mixed in 2% bovine serum albumin (BSA), was inserted into the cortical region of the brain, targeting the Kenyon cell cell body-rich region surrounding the calyx. The capillary tip was then withdrawn after dissolution of the dye crystal, a process typically completed within four seconds. Subsequently, the head was covered with fresh ringer solution and kept overnight in a dark and humid chamber to facilitate dye diffusion. The following day, the brain was dissected out, fixed in ZnFA, and immunostained with anti-SYNORF1 (3C11, Developmental Studies Hybridoma Bank, University of Iowa, IA, USA) following the same standard procedures described above and elsewhere [28,72]. Since the tract projections of Kenyon cells were scrutinized in whole-brain preparations, following immunolabeling, the brain underwent additional dehydration and clarification procedures. This involved immersing the brain in a series of baths containing increasing concentrations of glycerol (1%, 2%, 4% for 2 hours each, and 8%, 15%, 30%, 50%, 60%, 70%, 80% for 1 hour each) in 0.1 M Tris buffer with 1% DMSO, before complete dehydration with three consecutive washes in 100% ethanol (for 30 minutes each). The final clarification was achieved in Methyl Salicylate (M6752, MERCK Sigma Aldrich, MA, USA), which also served as both a storage and mounting medium.

### Neurotransmitters and Fasciclin-II

To assess MBON and DAN anatomy and examine lobe anatomy more closely, we used a selection of widely verified antibodies targeting conserved enzymes involved in neurotransmitter synthesis. Focus was put on those that were relevant for lobe anatomy, labelling whole cells to allow more accurate quantification. We emphasized understanding anatomy labelled by the antibodies rather than expression of the targeted proteins.

First, we used an antibody targeting GAD (Glutamic Acid Decarboxylase, #G1563, MERCK, Germany) which catalyses the conversion of L-glutamate to γ-aminobutyric acid (GABA). GABA is a prominent neurotransmitter and one of three found in MBONs [14]. There is a strong overlap between GABA and GAD antibodies as reported elsewhere [73], and western blot analysis identified bands consistent with GAD subunits [74]. Additionally, conserved labelling across arthropods has been identified [75], while GABA itself is highly conserved [76] giving confidence that GAD is an accurate marker for GABA-ergic neurons. Background staining common to GAD antibody labelling is easily identified and distinguishable from cell bodies, neurites and branching labelled by GAD.

Second, we used an antibody targeting TH (Tyrosine Hydroxylase, AB152, Merck Millipore, MA, USA) which synthesizes dopamine, the neurotransmitter expressed in dopaminergic neurons that carry valence signals related to reward or punishment. As it is a widely used and verified antibody, we were confident in its use to identify dopaminergic neurons [77–80]. Third, we used an antibody targeting serotonin or 5-HT (5-Hydroxytryptamin, #20080, Immunostar, Hudson, WI, USA) to identify further subdivisions independent of specific MBON or DAN labelling. The 5-HT antibody is widely used and verified across insects and other taxa, hence we were confident in its use to identify substructures (e.g. [81]).

We also used an antibody against Fasciclin-II (2F5, Developmental Studies Hybridoma Bank, University of Iowa, IA, USA) previously used in *Drosophila melanogaster* to reveal additional detail in the lobes, as it was reported to stain Kenyon cell populations and lobe divisions to differing intensities and play a role in neuronal guidance [34,82,83]. Our assessment shows a strong deviation from patterns in *D. melanogaster*, particularly by the strong labelling of the Y lobe that does not exist outside of Lepidoptera.

We also tested a Trio antibody (S1 Table), as Trio also labelled different lobe divisions as well as playing a role in axonal patterning [34,84] and easily accessible, but staining was unsuccessful (see tested conditions in S1 Table). We tested additional antibodies targeting other cross-reactivity of neurotransmitters reactive in other species, and report these tests and test conditions where they were unsuccessful in S1 Table for the benefit of the wider community.

### Imaging and image analysis

Imaging was performed with a Leica SP8 (Leica Microsystems) and a 20X air objective (20X HC PL APO CS2, NA = 0.75). We used a 65 mW Ar laser, a 20 mW DPSS yellow laser and a 50 mW 405 nm diode laser to excite Cyanine-2 linked signal, Cyanine-3 linked signal and DAPI, respectively. We used Hybrid detectors for Cyanine-2/3 and a PMT detector for DAPI. We aimed to scan a slightly larger view than one hemisphere at the time, guaranteeing a full view of the hemisphere in every slice. We used line averaging of 2-3, while using bidirectional scanning at 600 Hz with a resolution of at least 1024x1024. To calculate pinhole sizes we used either the average of all emission maxima in use or the smallest emission maximum and set AU = 1. We then used the system optimised z-slice size of 0.54 μm. We used linear Z compensation where necessary. Using the edge we cut into the agarose gel before slicing, we identified the same hemisphere consistently.

To generate accurate 3D segmentations, we corrected for axial aberrations due to refraction index differences between air and the mounting medium of 80% glycerol [85]. To determine a correction value for the z-axis, we used fluorescent beads with the fixed size of 14.6 μm (FocalCheck Microspheres, F7235, ThermoFisher, MA, USA) to compare their actual size to the imaged size using the 20x air objective. For this, we diluted these water-immersed beads for the end dilution of 80% glycerol. We then mounted them, and took two stacks, one with a system-optimised z slice value of 0.69 μm and one-over-sampled stack with the z slice of 0.2 μm. We then determined the beginning and end of the beads with the orthogonal views available in Fiji such that a continuous round surface was kept, took an appropriate substack and determined the stack size, and calculated the discrepancy between actual and measured size. We repeated this for 3 beads per stack type and calculated a mean (and standard deviation) of 11.97 ± 0.48 μm, to determine a correction value of 1.22.

### Image processing, 3D segmentation and annotation

We used Fiji 1.54d [86] and the included standard tools to modify brightness/contrast, orientation in all axes and other standard procedures. To generate a merged whole brain picture from brain slices, we reoriented them in their rotational axis and z-orientation (*reverse, rotate, flip horizontally*), we removed any unnecessary slices using *Substack*, and then used *Concatenate* to stitch all z-stacks from one brain together, before separating the channels. To exactly align the concatenated z-stack in both X and Y, we used Amira 3D 2021.1 (ThermoFisherScientific, MA, USA). For this, we first used the correct voxel size for each stack, with the z-value multiplied by the correction value of 1.22. We then used the module *Align Slices* in the red-green view. Here, we went to the slice where stack 1 ends, showing colour-indicated mismatches. By shifting the image and dragging it in X/Y, these mismatches were removed. Repeating this for all borders where one stack led into the other, an alignment was achieved. Resampling then produced the final aligned stack of the first channel. Using *Align slices* on the second channel, we then used the first aligned channel as reference, assuring that we had identical changes throughout both channels and recorded signals. Both channels were saved to proceed with 3D segmentations.

We segmented the brain and lobe parts using the *labelfield* module and manual segmentation in approximately every 5-10 slices (2.5-5.5 μm), using interpolation afterwards. We verified the precision of interpolation and our segmentation using all views available. We then smoothed the selection, as well as the segmentation later on with size 8 in all axes. Using Material Statistics we extracted volumes of each subdivision segmented. To generate a surface view we used constrained smoothing at a level of 3.

Note that we segmented differing numbers of subdivisions between species as greater numbers of lobe subdivisions were detectable in *Dryas iulia* and *Dione juno* in contrast to the *Heliconius sp*. While we used these as is for anatomical descriptions (Fig 1), for statistical analysis, we added values for both α’ lobes and β’ lobes as well as for γ’ and γ and γ lobelets that we were able to consistently identify in *Dryas iulia* and *Dione juno* but not in *Heliconius sp*. This way, we captured as much detail as possible while also allowing for comparative statistical analyses.

We used the nomenclature established in Ito et al. [20] where applicable, and adhered to the nomenclature of lobe structures established by Heisenberg [18] and used in previous work [14,19]. We always oriented the optic lobe to the left and midline to the right. For ease of comprehension, we treated the orientation of the mushroom body as if it were on the body axis from anterior to posterior, i.e. the lobes were the most anterior and the calyces the most posterior. This made more complex descriptions of locations easier. We used Inkscape (https://inkscape.org) to generate all figures.

## Statistical analysis

All statistical analysis performed in this paper can be found in the S1 Script. Statistical packages, the data set and full results are reported in S2 Table. We report the general procedure here and link to specific detail to the Supporting Files.

To test for species- and clade-specific effects of KC expansion, we first performed several GLMMs (General Linear Mixed Models) and standard LMs (linear models). Importantly, as we wanted to test whether the different lobe divisions expanded all together or independently from each other, we structured the dataset by having volume as dependent variable and structure identity as a test independent variable, alongside species, clade identity, sex and age as additional test variables and identifier as a control random factor. Model diagnostics were performed for multiple regressions using the package *ggResidpanel* and for GLMMs using the package *DHARMa*. In both cases, we tested for normal distribution of residuals, heteroscedasticity, and residual distribution across the independent variables. Test plots can be recreated using the S1 Script. All model results described have been selected and diagnosed in such a way. Only the selected model of each set of nested models was reported and interpreted. Procedures and all models can be found in the S1 Script.

To test sex differences, we designed a set of nested models including or excluding sex. From these, we selected the model with the lowest AIC value (package *bbmle*) and simplest terms. Results of these sex differences are reported in detail in S1 text. To test age differences, we generated a dataset of five 9-10 day old *Dryas iulia* and *Heliconius erato* that we combined with day 1 individuals that were used in the other analyses. We used an appropriate model and compared it to a null model to test for these effects, and subsequently subset the data to test for structure-specific effects using standard linear models (S2 Table). In the main set of models where we tested for species differences, we also wanted to identify whether the effects of clade (*Heliconius* vs sister group) were significantly different from single species effects. For this, we compared the two models using ANOVA. To perform posthoc effects of species on each structure, we used the package *eemeans* (S2 Table). To retrieve confidence intervals of R^2^ from these posthoc tests, we used a bootstrapping approach with 1,000 iterations.

Principal Component Analyses (PCA) were performed using standard procedures (Fig 2d). *smatr* analyses were performed by generating a robust standardised major axis (SMA) regression, including a Huber’s M estimation that tested for slope and elevation differences (Fig 2E, S4 Fig)[41,87]. In these analyses we additionally correct for multiple testing to identify pair-wise species effects (S2 Table). Tests for cell counting (Fig 3) were performed with standard linear models.

P-value adjustments were performed using the Benjamini-Hochberg corrections, and applied in the case of the single structure species (Fig 2B) and age models (S3 Fig), by multiplication of p values by six (as six structures were used). P-value adjustments in the pairwise *smatr* analysis (S4 Fig) were performed by 10, as each subdivision was compared 10 times.

## Author contributions

SHM conceived, supervised and administered the study, and obtained funding. MSF and SHM designed the final methodology study. AC established experimental techniques and performed calyx injections. MSF and TL performed all other labelling experiments. MSF imaged, 3D segmented and analysed the data. MSF and SHM interpreted the data, and MSF wrote the first draft of the manuscript, prepared the data visualisations and curated the data, with input and editing from SHM. All authors read and approved the final version of this work.

## Acknowledgements

We thank Francesco Cicconardi for help with identifying orthologous antigens to the antibodies we used, Amaia Alcalde for providing a larval specimen as well as help with larval rearing, Elizabeth Hodge for re-imaging some injected brains, and Katherine Pearson-Bunt having been involved in early iterations of this work. We acknowledge help by Philip Steinhoff regarding alignment of z stacks in Amira as well as Elizabeth Martin-Silverstone for technical assistance. We gratefully acknowledge the Wolfson Bioimaging Facility, particularly Katy Jepson and Dominic Alibhai, for their support and assistance in this work. This project was instigated under a NERC Independent Research Fellowship NE/N014936/1 and ERC Starter Grant 758508, and completed under HFSP Project Grant RGP0029/2022 to SHM. MSF was supported by a Walter-Benjamin Fellowship from the Deutsche Forschungsgemeinschaft (FA 1818/1-1).

