## Supporting Figures and Text for "Mosaic evolution of a learning and memory circuit in Heliconiini butterflies"

**Title:**

**Affiliation:** [a] Evolution of Brains and Behaviour lab, School of Biological Sciences, University of Bristol, Bristol, United Kingdom; [b] Evolution, Genomes, Behaviour and Ecology Lab, Centre national de la recherche scientifique (CNRS), Université Paris-Saclay, France.

**Correspondence**:

MSF:

SHM:

**ORCiD**:

MSF https://orcid.org/0000-0003-2418-3203

AC: https://orcid.org/0000-0002-5528-7331

SHM: https://orcid.org/0000-0002-5474-5695

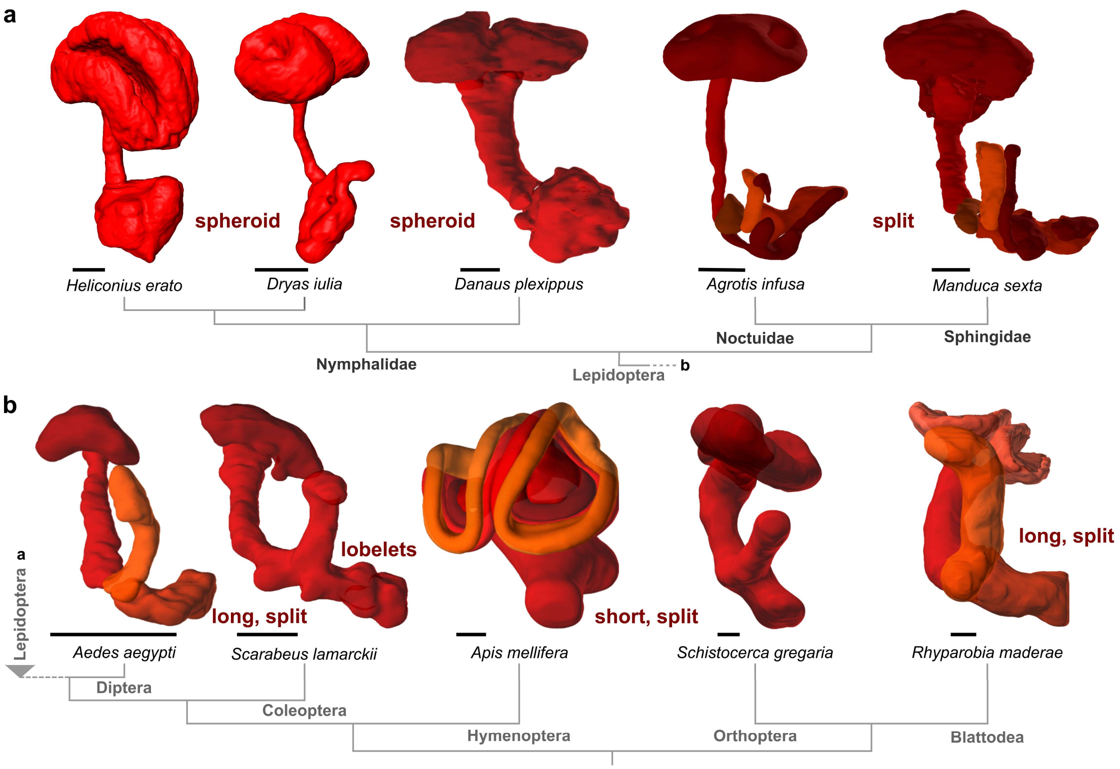


**S1 Fig: Morphological variation in mushroom bodies, with emphasis on the lobes in select species of Lepidoptera (a) and representatives of other insect orders (b) (not to scale)**. Generally, lobe shape differs in the degree of separation into a vertical and medial lobe, resulting in a lack there-of and hence spheroid structures inside Nymphalidae, or long or short lobes in other orders. Sometimes, very specific lobelets are visible. Phylogenetic relationships are indicated by a cladogram, based on standard phylogenies [91]. With the exception of the Heliconiini, 3D models of all species were extracted from insectbraindb.org. Importantly, the 3D models differ in the exactness of the lobes displayed here. Note large differences in absolute size (all scale bars are 100 μm). Y tract in Lepidoptera not shown due to lack of consistent availability of data.

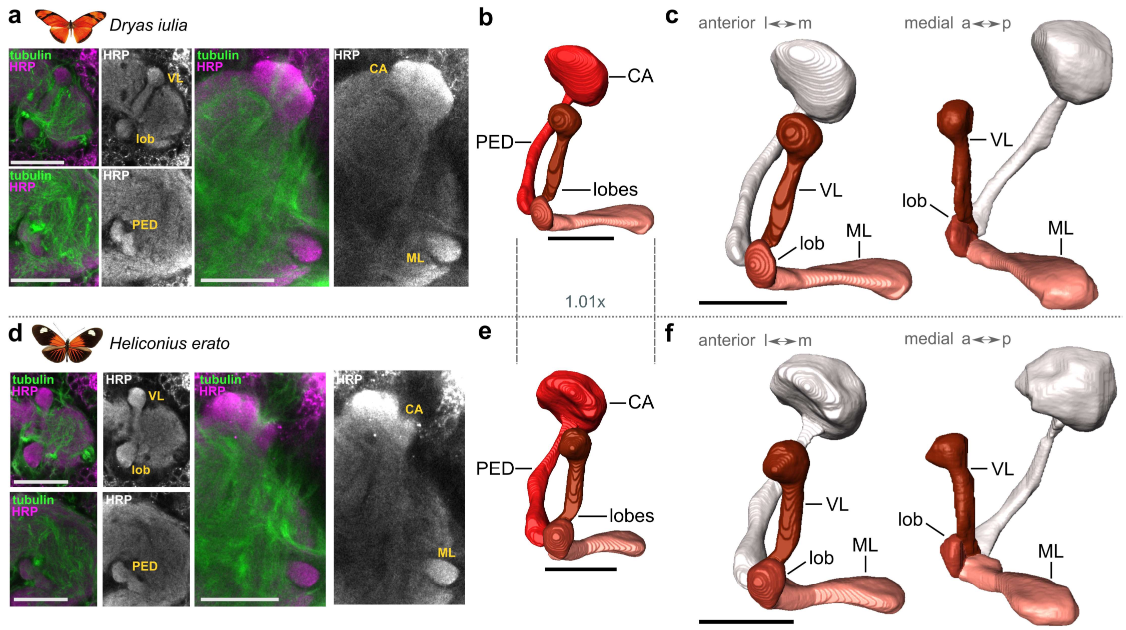


**S2 Fig: Large degrees of conservation in lobe anatomy between L2 larvae of a non-pollen feeder *Dryas iulia* (upper row) and a pollen feeder *Heliconius erato* (lower row)**. **a**/**d**. Singe plane sections of stainings in *Dryas iulia* (**a**) and *Heliconius erato* (**d**) that were used to determine the different subdivisions, including denotations of structures. **b**/**e**. 3D segmentations of the larval mushroom bodies show a lack of absolute size divergence. **c**/**f.** Shown are visible subdivisions in the lobes in both species in two different positions. The lobes are mainly divided in a medial (ML) and vertical section (VL), with a bulbous structure, a lobelet (lob) at the split point of the peduncle (PED), and a posterior calyx (CA). All scale bars are 50 µm.

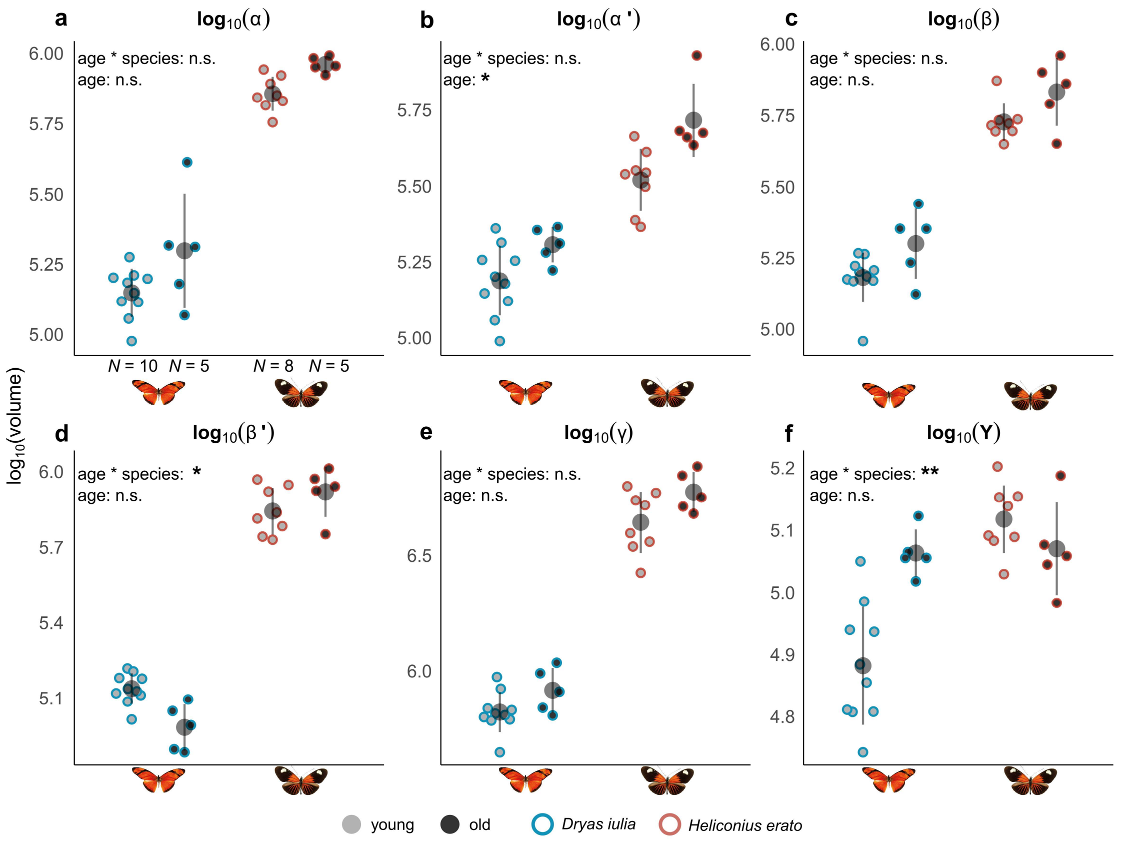


**S3 Fig: Analysis of age effects across 9-10 day old and 1 day old *Dryas iulia* and *Heliconius erato* across each subdivision** (**a-f**). Indicated are results of a general linear model with an interaction of age and species.

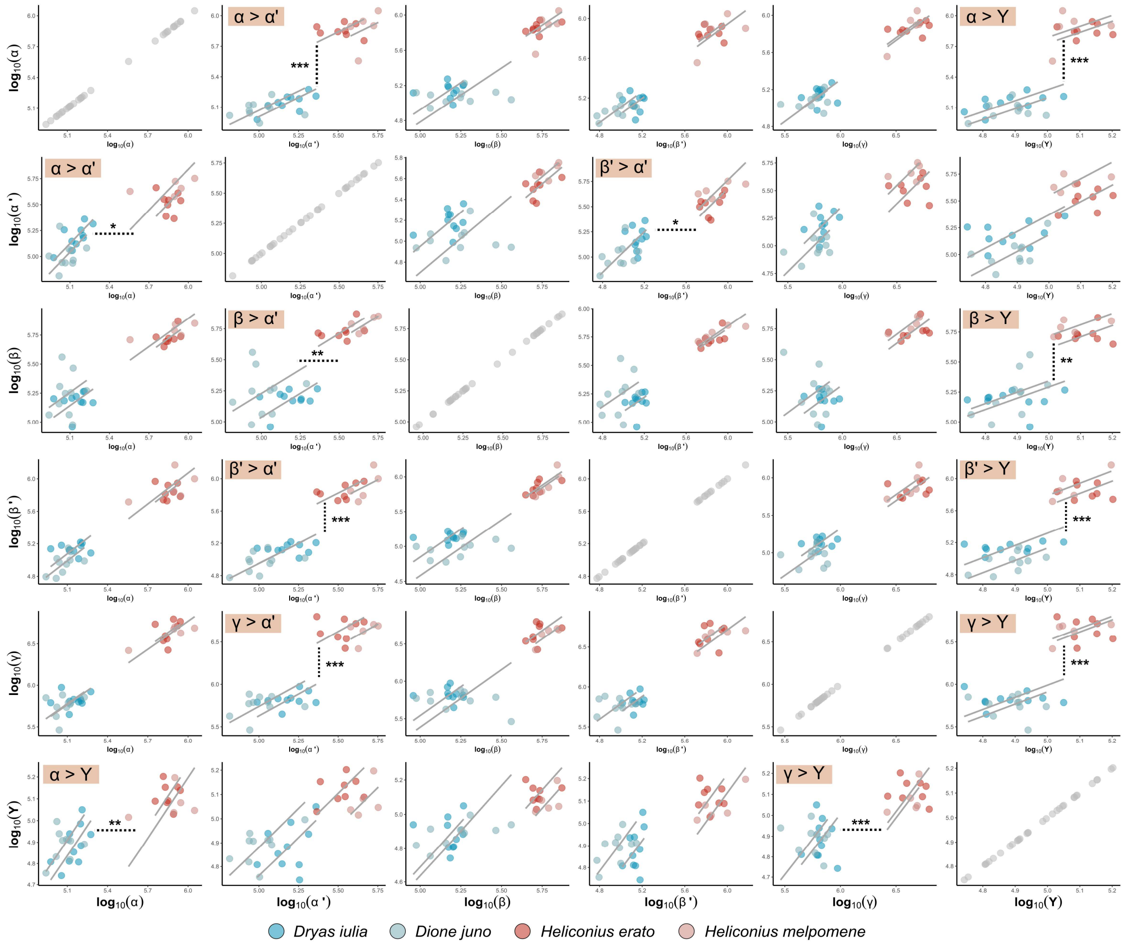


**S4 Fig: Comprehensive pair-wise analysis of scaling relationships between all combinations of lobe subdivisions by *smatr* analysis**. Each plot includes indications of significance of elevation differences between *Heliconius* and the two outgroup species (species-specific differences are reported in the data file). Where broad elevation differences were significant, expansion patterns between each two divisions are indicated in the upper-left corner. Colour-coding for species found below.

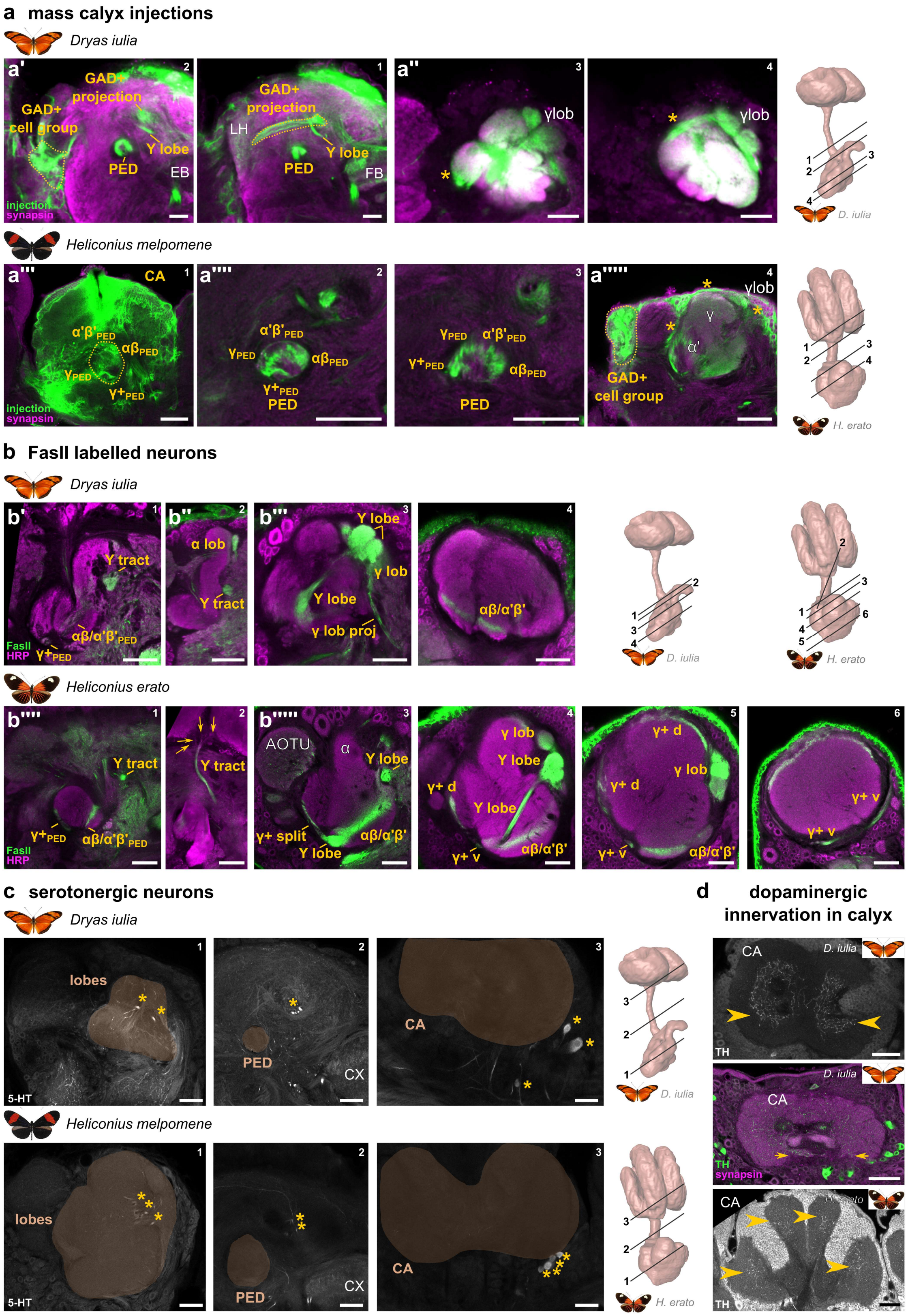


**S5 Fig: Additional staining techniques reveal necessary detail about the Heliconiini mushroom body lobes.**

**a.** Mass injections of Dextran into the calyx reveals: i) a GAD-positive cell group of GABA-ergic feedback neurons (as in Figure 3) and its bifurcating projection into the calyces, its labelling through calyx injections confirming this anatomy (a',a'''''); ii) three inputs into the mushroom body lobes, through the peduncle, the Y tract and the GAD-positive projections labelled here (a'); iii) confirmation that the γ+ division of the peduncle indeed innervates the γ lobelets (a'',a'''''); iv) rotation of the Kenyon cell populations throughout the brain from calyx into lobes (a''', a''''). **b.** Analysis of FasII labelled neurons reveal specific subpopulations of Kenyon cells, particularly those which make up the Y tract and innervate the Y lobe. Moreover, the γ+ division of the peduncle is labelled specifically, as well as a part of neurons in the border between αβ and α'β' lobes. There are indications of clade differences between *Heliconius* and outgroup species. **c.** Identification of a conserved group of serotonergic neurons through an antibody targeting 5-hydroxytryptamine (5-HT) that innervate the lobes through its dorsal point and have their soma near the calyx. **d.** Shown is a detail revealed through Tyrosine Hydroxylase (TH) antibodies of dopaminergic innervation in the calyx, where specific edges were labelled, particularly in the outgroup genera, and to a lesser, but still visible extent in *Heliconius* as well. All scale bars are 50 µm except for a'''-a''''' (100 µm). 3D segmentations indicated the location of each shown image, identified by number in the upper right corner. 3D segmentations are from identical sources than in Fig 1. Abbreviations: PED peduncle, EB ellipsoid body, LH lateral horn, FB fan-shaped body, lob lobelet, proj projection, d dorsal, v ventral, CA calyx.

**Anatomical description of lobe anatomy**

To provide deeper insights into the anatomy of the Heliconiini mushroom body lobes, we provide here a detailed description.

The α/β portion of the peduncle split posteriorly into a prominent vertical lobe, making up a large portion of a vertical component particularly in *Dryas*, and reaching into rather posterior locations compared to the other divisions. Medially, we identified a spheroid β lobe in *Dryas* while it was a flat layer very ventrally in *Heliconius* species. The α lobe was less prominent in *Heliconius* as it is covered anteriorly by a massive γ lobe. The next division, α'/β', is split further medially into a vertical α' lobe that is nestled between the more dorsal α and the more ventral γ and β' lobes. β' lobes in *Dryas* are made up by two subregions, most ventrally to all other sections, while in *Heliconius* species they are in between the γ and β lobe and one mass. The γ lobe then makes up a massive portion of the lobes in both species. We identified a separate γ' lobe in *Dryas*, which we were not able to identify in *Heliconius*. Note that we used FasII labelling as a suitable way of distinguishing the border between α/β and α'/β' as it labelled a thin layer of the first (S5 Fig B).

We also described several additional subportions, particularly a very distinct portion of the peduncle that only seemed to diminish very anteriorly, termed here 'γ+'. Through Dextran injections into the calyx, we were able to confirm that this peduncle division supplies the γ lobelets, that surround the medial portion of the Y lobe (Fig 1e/g, S5 Fig a''/a'''') in both clades. In *Dryas*, we also identified slices in the same division, e.g. α' and β', indicative of further Kenyon cell types not identified through our methods (note molecular descriptions of Hymenopteran Kenyon cells describe [88]10 cell types [88]). Additionally, we identified a lobelet next to the very posterior tip of the α lobe, termed α lobelet. This was particularly visible in *Dryas*, less so in *Heliconius*, but its shifted position and identity was confirmed through FasII stainings (S5 Fig).

The Y lobe is a prominent feature of Lepidopteran brains that is supplied by Kenyon cells that run through their own distinct tract, the Y tract, and not the peduncle, which is supplied by four branching neurites originating from cells that are ventro-anterior to the main grouping of cells in the calyx (Fig S5b'''' inset 2)[19,89]. The Y tract projects first sharply anterior, then ventrally, then anteriorly again and ends in the medial side of the lobes approximately at middle height. This conflation point splits and we see a very conserved picture of two medial lobelets and 1 lateral lobelet connected by a tract running through the lobes (dividing the portions into a vertical and medial identity).

Note that there is some contention in the literature regarding the identity of α and α' lobes. We see in some publications [23] that the posteriorly and often dorsal reaching section of the vertical lobe has not been identified as α, while it has in others [13,19]. We see quite close resemblance of the vertical lobe, particularly from a lateral perspective and through its reach into the posterior part of the brain, with *Drosophila* and other species [13,14] and have thus identified the posterior-dorsal portion as α and not α'.

We also wanted to explore the anatomy of the larval mushroom body lobes to examine a) when the large adult differences are developed and b) whether there are any differences in lobe anatomy during the larval stage between the clades (S2 Fig). For this we made tubulin/HRP stainings in early L2 stage larva brains and segmented their lobes. The larval lobes at this stage and across larval development (Hebberecht pers. obs.) are mainly separated into a medial and vertical lobe (ML and VL, respectively), as rather long thin tubes. There is an additional lobelet very close to the peduncle, not resembling a spur like in adults of other insect species [13,37], but it is very much anterior to the peduncle split (S2 Fig C/F). The vertical section strongly resembles the α lobe, while the shape of the medial lobe has no equal in adult anatomy, but stretches relatively far into posterior, while the VL stays anterior (in contrast to the adult α lobe it resembles). As such, the larval mushroom body lobe is very similar to the basic structure of the *Drosophila* larval mushroom bodies [39,40]. Considering there are obvious morphological and behavioural differences between *Drosophila* and Heliconiini larvae – such as multi-joint legs or gregariousness [90] – it is surprising not to find obvious anatomical differences in the mushroom bodies. It is possible that a larval central complex, present and functional in Heliconiini (Farnworth, pers. obs.) but absent in *Drosophila*, facilitates some of the larval differences. However, it would be interesting to explore the mushroom bodies and Kenyon cell populations more in Heliconiini to potentially relate differences in Kenyon cell types or distribution of DANs or MBONs to behavioural adaptations. In fact, in the *Drosophila* larvae all Kenyon cell axons in the lobes are γ Kenyon cells as they are born first [39]. It is so far unclear as to which types of Kenyon cells the cells in the Heliconiini larval lobes belong to and whether the temporal cascade is conserved. Importantly, we see no obvious differences between species at this larval stage, which also implies that all differences present in the adult must have arisen during late larval and pupal development.

**Additional statistical results**

We had two additional independent effects we tested, complimentary to the main effect of species identity: age and sex.

First, as we had limited availability of data on sex in the dataset (24 out 43 individuals, i.e. 56%), we tested sex effects in this subset of 24 individuals. After model selection (see S1 Script), we ran model 1 (see S2 Table) in the subset of 24 individuals, revealing no significant relationship between lobe structure volumes and sex upon comparison to the corresponding null model. We concluded that there are no obvious sex differences in the size of each lobe structure and have ignored sex in all subsequent analyses.

We next wanted to test for age effects in lobe structure sizes in a sub-group of our data in the species *Dryas iulia* and *Heliconius erato* where we had data for 9-10 day old butterflies (classed “old”, 5 individuals per species) and freshly eclosed day 0 butterflies (classed “young”, 10 *Dryas iulia*, 8 *Heliconius erato*). Species specific age effects in learning effects and mushroom body anatomy have been reported previously [29], hence why we tested additionally for age-specific species effects across the sizes of the different lobe structures. We identified that upon comparison to the null model, there were species-dependent age effects on lobe structure size (i.e. the interaction term age*species being significant; $X^2$ = 45.353, $P < 0.001$) and age generally had an effect on structure sizes ($X^2$ = 61.466, $P < 0.001$). To explore these age effects on single structures further, we subsetted the data to test for species-dependent age effects in each structure separately (S3 Fig). These *post hoc* models identified significant species-dependent age effects in the β’ and Y lobes, where we had inverse trends in each species. Specifically, we identified smaller β’ in older *Dryas iulia*, but larger ones in *Heliconius erato,* and larger Y lobes in older *Dryas iulia*, but smaller ones in *Heliconius erato*. We also identified consistent age effects across both species in α’, where older individuals had slightly larger α’ lobes. Importantly, in contrast to other published work, these age effects were from laboratory-reared animals, potentially explaining differences in effect size and some rare but interesting species-specific age effects. Due to the small effects we were able to identify, we refrain from larger interpretation here, but hypothesise a potential mosaic response to aging across these lobe divisions.
